# The GTPase Activity of the Double FYVE Domain Containing Protein 1 (DFCP1) Regulates Lipid Droplet Metabolism

**DOI:** 10.1101/2022.02.09.479746

**Authors:** V.A. Ismail, T. Naismith, D.J. Kast

**Affiliations:** Dept. of Cell Biology and Physiology, Washington University School of Medicine, St. Louis MO

## Abstract

Lipid droplets (LDs) are transient lipid storage organelles that can be readily tapped to resupply cells with energy or lipid building blocks, and therefore play a central role in cellular metabolism. However, the molecular factors and underlying mechanisms that regulate the growth and degradation of LDs are poorly understood. It has emerged that LD metabolism is sensitive to the autophagy- and LD-associated protein Double FYVE Domain Containing Protein 1 (DFCP1), however, little is known about DFCP1’s roles in autophagy and LD metabolism. Here, we show that DFCP1 contains a novel GTPase domain that regulates LD size by controlling the assembly of DFCP1 onto LDs in response to changes in nutrient availability. Specifically, we show that DFCP1 accumulation on LDs is independent of PI3P-binding, but requires a combination of the ER-binding domain and a unique GTPase domain. This novel GTPase domain possesses a low basal GTP turnover rate and has the ability to dimerize. Furthermore, mutations in the DFCP1 that either impact GTP hydrolysis or dimerization, result in changes in the accumulation of DFCP1 on LDs, as well as in changes in LD density and size. Importantly, the magnitude of these changes depends on the nutritional status of the cell. Collectively, our findings indicate that DFCP1 is a GTP-dependent metabolic sensor capable of modulating cellular storage of free fatty acids.

## Background

Lipid droplets (LDs) are conserved transient lipid storage depots that can provide lipids for the repair and biogenesis of membranous organelles, as well as serving as a source of energy during times of nutrient stress. LDs accumulate in response to excessive dietary fatty acids (FAs) and can be broken down in response to meet the energy demands of the cell or the entire organism. Contacts between the endoplasmic reticulum (ER) and LDs are essential in allowing for cellular lipid metabolism by providing avenues of lipid traffic into and out of the LD. Failure to store or catabolize lipids from LDs results in a host of metabolic diseases including lipodystrophies, obesity, insulin resistance, diabetes, NAFLD, and atherosclerosis (Krahmer et al., 2013). However, the molecular factors that regulate LD growth and degradation remain unknown.

DFCP1 (also referred to by the gene name ZFYVE1) has historically been used as a marker for the early steps of autophagy, where it is known to accumulate at phosphoinositol-3-phosphate (PI3P)-rich autophagosome precursor sites on the ER upon the induction of macroautophagy (Axe et al., 2008). This localization is due to a unique combination of structural elements that include an ER localization domain and a tandem pair of PI3P-binding FYVE domains (Ridley et al., 2001). At these sites, DFCP1 was shown to mediate ring-like extensions of the ER that precede the accumulation of the autophagosome marker, LC3-II. However, in spite of this specific localization, DFCP1 appears to be dispensable for macroautophagy (Axe et al., 2008).

More recently, DFCP1 was shown to localize to lipid droplets (LDs) using a proximity assay to probe for potential interactors of the rate-limiting lipolytic enzyme, ATGL, on the LD surface (Bersuker et al., 2018). It has since been shown that overexpressed GFP-DFCP1 accumulates on LDs in cells, and this overexpression increases their size (Li et al., 2019), whereas knockdown of DFCP1 was shown to do the opposite – decreasing LD size in exchange for an increase in the number of LDs. Interestingly, the function of DFCP1 on LDs may be indirect, in that it was proposed to assist in the recruitment of BSCL2, as well as Rab18 and ZW10 (Li et al., 2019), which are two components of the NRZ ER-LD contact site complex (Xu et al., 2018). However, it should be noted that studies on the Rab18 interactome (Kiss et al., 2019) and/or the purified ER-LD tethering NRZ complex (Hirose et al., 2004) have not identified DFCP1 as a potential interactor. Thus, it is not clear whether DFCP1 can directly modulate LD metabolism or whether its potential association with autophagy can be integrated with its role on LDs. For this purpose, we studied DFCP1’s localization to LDs under conditions that induce macroautophagy and to determine the molecular factors that allow DFCP1 to switch between LDs and/or autophagosome precursor sites.

Here, we show that DFCP1 is an important regulator of LD metabolism. In particular, we show DFCP1 impedes LD formation and FA uptake, and that ablating DFCP1 markedly enhances cellular LD capacity. Mechanistically, this ability to block FA storage depends on a previously unreported GTPase domain, and while enzymatically similar to Ras-type GTPase, the DFCP1 GTPase domain bears little amino acid sequence similarity with known GTPases. Importantly, mutations that alter DFCP1’s GTPase activity lead to aberrant localization and accumulation of DFCP1 on LDs. Overall, we have re-classified the role of this established marker of autophagy, showing that GTP-dependent localization of DFCP1 to LDs directly impairs excess cellular storage of FAs.

## Results

### DFCP1 associates with LDs during macroautophagy

The intracellular localization of DFCP1 is known to be sensitive to nutrient stress (Axe et al., 2008). Under normal growth conditions, DFCP1 localizes to the ER and/or the Golgi in U2OS cells (**Figure S1A**). However, when these cells are depleted of nutrients by replacing the growth media with EBSS (hereafter referred to as starvation), they undergo a process called macroautophagy, and, as a consequence, DFCP1 accumulates on LC3-positive subdomains of the ER that are assumed to serve as autophagosome assembly sites (**Figure S1A**). Curiously, this localization appears to be coincidental, since knockdown of DFCP1 does not affect autophagosome biogenesis, as indicated by the processing of the cytosolic LC3-I into the autophagosome bound LC3-II (**Figure S1B**).

More recently, it has been shown that when cells are presented with a surplus of fatty acids (FAs), DFCP1 preferentially localizes to lipid droplets (LDs) (Bersuker et al., 2018; Gao et al., 2019; Li et al., 2019), however, it is not known if this localization is sensitive to starvation. Under fed conditions, GFP-DFCP1 localizes strongly to the periphery of LDs in osteosarcoma (U2OS) cells stimulated to form more LDs by supplementing the growth media with 200 μM exogenous oleic acid (OA) (**Figures 1A-C**). Inducing macroautophagy in the same cells leads to both a loss of DFCP1 localization with LDs (**Figure 1B**) as well as the amount of DFCP1 accumulated on LDs (**Figure 1C**). Importantly, DFCP1 reorganizes from a symmetric arrangement enveloping an LD to an asymmetric accumulation on one side of the LD (**Figures 1A and 1D**) that is typically seen at the junction between neighboring LDs or the ER-LD contact site. This starvation-dependent reorganization on LDs persists even when these OA-treated cells are starved and/or treated with PI3K inhibitor Wortmannin (**Figures 1A and 1B**), suggesting that PI3P binding by DFCP1 is dispensable for its localization to LDs. In support of these observations, LDs purified from OA-stimulated U2OS cells have less endogenous DFCP1 bound when cells are starved for 4 h prior to purification. Importantly, this accumulation on LDs is independent of the ER, as the ER marker calreticulin was completely absent in the purified LD fraction **(Figures 1E and S1C)**. Furthermore, GFP-DFCP1 forms intense shell-like accumulations on LDs purified from fed cells, and this is diminished in those LDs purified from starved cells (**Figure 1F**). These findings strongly suggest that DFCP1 can directly interact with LDs and that this interaction is attenuated in starved cells.

**Figure 1:**
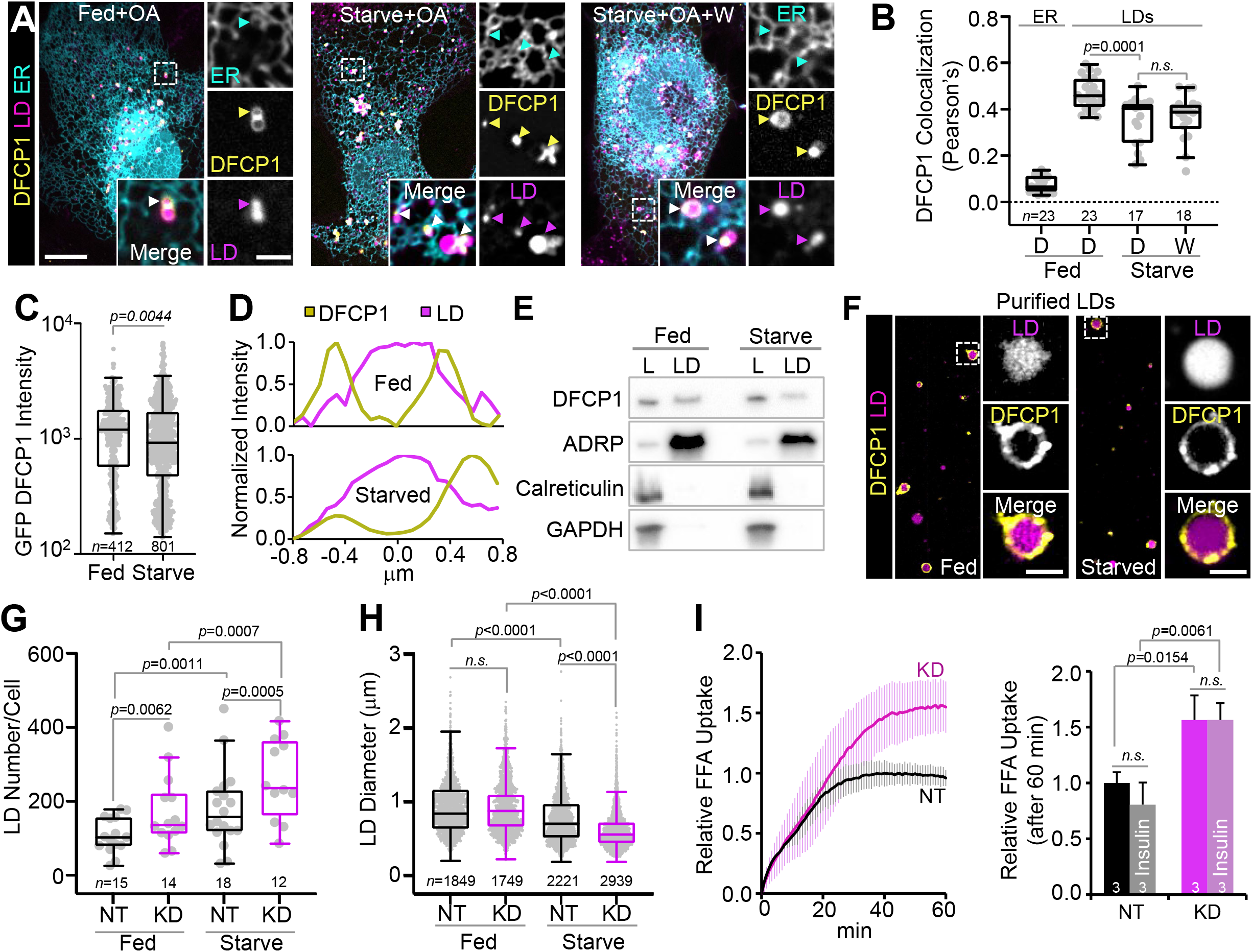
DFCP1 Regulates LD Metabolism. **(A)** Live cell confocal images of U2OS cells expressing BFP-DFCP1 and mCherry-Sec61β. Cells were treated with 200 μM oleic acid (OA) for 4 h to stimulate LD formation (OA-stimulated) prior to 1 h incubations in growth (left), starvation media (middle), or starvation media supplemented with 1 μM Wortmannin for 30 min (right). LipoTox Green was used to mark LDs. **(B)** Bb)Colocalization (Pearson’s correlation coefficient) of BFP-DFCP1 with the ER and LDs from fed and starved U2OS cells treated with either DMSO (D) or 1 μM Wortmannin for 30 min (W). **(C)** Accumulation of GFP-DFCP1 (Log of fluorescence intensity) on LDs in fed and starved, OA-treated U2OS cells. **(D)** Representative line scans across LDs formed fed (top) and starved (bottom) OA-stimulated U2OS cells shown quantified in **Figure 1C**, expressing BFP-DFCP1 (gold) and GFP-Sec61β and stained with LipoTox Deep Red (LipoTox DR) (magenta). **(E)** Western blots of purified LDs isolated from fed and starved OA-stimulated U2OS cells showing the total cell lysate (L) and the purified LD fraction (LD). **(F)** Confocal images of purified LDs isolated from fed OA-stimulated U2OS cells expressing GFP-DFCP1 and stained with LipoTox DR. **(G,H)** Number and diameter distributions of LDs quantified from images of control non-targeting siRNA treated (NT) and DFCP1 siRNA (KD) OA-stimulated Hep3B cells expressing LifeAct-mTagBFP2 and GFP. LD diameter was measured in the plane of a confocal Z-stack where a given LD’s diameter was the greatest. LDs were visualized using LipoTox DR. **(I)** FFA uptake measured in NT and KD Hep3B cells cultured under normal growth conditions. Lines represent mean ±SD for 3 replicates in 1 experiment. The scale bars in whole-cell and inset images represent 10 and 2 μm, respectively. The statistical significance of the measurements was determined using the Mann–Whitney U-test on the indicated number of observations (indicated in each figure panel) recorded from two independent transfections. Exact p-values are reported with exception to p>0.05, which are considered to be nonsignificant (n.s.) See also **Figure S1**.

Localization of DFCP1 to LDs has previously been suggested to regulate LD growth but not biogenesis, since loss of DFCP1 drives the accumulation of LDs that are smaller on average compared to those found in fed WT U2OS cells (Li et al., 2019). We also observed a similar trend in hepatocellular carcinoma (Hep3B) cells, except in these cells, we found that RNAi-based depletion of DFCP1 (hereafter referred to as DFCP1 KD) led to an increase in LD number when compared to scrambled-treated Hep3B cells, whereas the LD size distribution (as determined by the maximum diameter of each and every LD in the cell) was unaffected **(Figures 1G and 1H)**. Interestingly, this implies that DFCP1 KD induces a net increase in lipid storage capacity, which correlates well with a ∼50% increase in insulin-dependent and insulin-independent palmitate uptake (**Figure 1I**), and intracellular FFA content (**Figure S1D**).

DFCP1’s role on LDs during nutrient stress is also of interest as LD growth and degradation are intertwined processes. Upon starvation, cells will readily tap into their lipid stores to help replenish ATP levels and other amino acid building blocks. This catabolism of LDs in hepatic cells requires two potentially sequential mechanisms involving lipolytic breakdown of large LDs by LD-associated lipases into small LDs, which can then be efficiently cleared by the autophagy-lysosomal pathway, in a process known as lipophagy (Kaushik and Cuervo, 2015; Schott et al., 2019; Schulze et al., 2017). Interestingly, the clearance of LDs can lead to a surplus of FAs, which have been shown to be repackaged into small nascent LDs (Nguyen et al., 2017). Thus, starvation induced breakdown of LDs causes a net increase in the number of LDs at the expense of LD size, as exemplified in transfected Hep3B cells (**Figure S4E**). Interestingly, this starvation-induced accumulation is further exacerbated by DFCP1 KD, which leads to LDs that are significantly more abundant and smaller than those found in either fed control, fed KD cells or starved control Hep3B cells (**Figure 1G and 1H**). Altogether, this suggests that DFCP1 helps to protect LDs from starvation induced catabolism.

### DFCP1 localizes to LDs using two distinct structural domains

In the presence of LDs, DFCP1 is less sensitive to this starvation-induced translocation, which helps to protect LDs from catabolism. This suggests that additional intramoleclular factors counteract PI3P-mediated transmigration of DFCP1 to sites of autophagosome formation. The localization of DFCP1 to either the ER or LDs is known to require the ER-binding motif (ERB) (Axe et al., 2008; Gao et al., 2019; Li et al., 2019), and starvation-induced translocation to PI3P-rich regions is known to also require the C-terminal tandem FYVE domains (Axe et al., 2008; Cheung et al., 2001). To identify these additional domains that may contribute to DFCP1’s localization to LDs, we transiently expressed GFP-DFCP1 truncations (**Figures 2A, S2A and S2B**) in fed and starved OA-treated U2OS cells in order to identify the domains that could contribute to DFCP1’s localization to LDs. In addition to the C-terminal FYVE domains (FYVE), and the ERB, DFCP1 has an N-terminal domain that bears some similarity to a zinc finger domain (N) and a folded domain containing a canonical P-loop sequence (P-loop) (**Figure 2A**). Among all the GFP-constructs tested, those constructs containing the ERB (1-777, 112-553, 415-553, 415-777, 112-777, 1-553) show greater LD colocalization (as determined under both fed and starved conditions **(Figures 2C-F, S2D and S2E**), as determined by Pearson’s correlation coefficient (**Figures 2B and S2H**). Notably, this colocalization is stronger than that seen by the promiscuous ER marker, Sec61β, indicating that these constructs specifically associate with LDs, as opposed to residing on the ER that is in proximity to the LDs (**Figures 2B, S2C, and S2H**). Constructs lacking the ERB (**Figures S2F and S2G**) are completely cytosolic in both fed and starved cells, and therefore show no colocalization with LDs (**Figure S2H**). Importantly, this is not a result of excessive transient expression, as constructs lacking the ERB display more attenuated expression (**Figure S2B**), compared to those that do contain the ERB.

**Figure 2:**
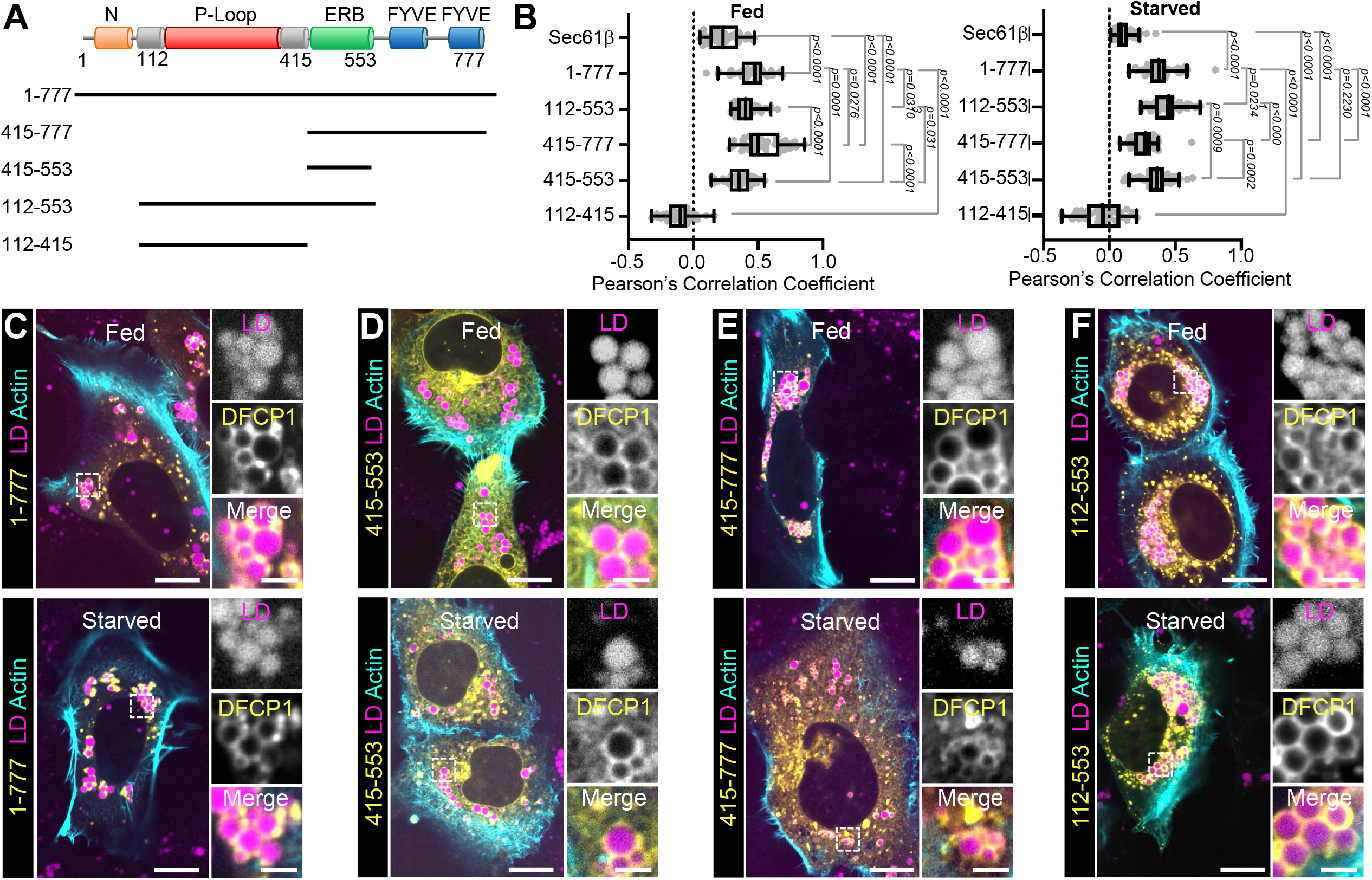
Domain Requirements for DFCP1 Localization to LDs. **(A)** Domain diagram of DFCP1 depicting the N-terminal domain (N), the GTPase domain (P-loop), the ER-binding domain (ERB), and the tandem FYVE domains (FYVE). GFP-tagged constructs used in this figure are depicted below the domain diagram. **(B)** The extent of colocalization (Pearson’s correlation coefficient) between GFP-DFCP1 and LDs from fed (left) and starved (right) cell populations depicted in **Figure 2C-F**. The extent of colocalization between GFP-Sec61β and LDs is also included as a reference. **(C-F)** Representative images of U2OS cells expressing LifeAct-mTagBFP2 and the GFP-DFCP1 truncations indicated in **A**. Prior to imaging, all cells were OA-stimulated for 20 h before incubating in either growth or starvation media for 18 h. LDs were visualized with LipoTox DR. The scale bars in whole-cell and inset images represent 10 and 2 μm, respectively. The statistical significance of the measurements was determined using the Mann–Whitney U-test. Exact *p*-values are reported with exception to those that are >0.05, which are considered to be nonsignificant (n.s.) See also **Figure S2**.

While the ERB is essential to localize DFCP1 to LDs, it does not, however, confer sensitivity to starvation. The ERB domain construct (415-553) localizes equally to both the ER and LDs, in both fed and starved cells (**Figures 2B and 2D**). By contrast, a construct that contains both the ERB and the FYVE domains (415-777) strongly translocates away from LDs in starved cells (**Figures 2E and 2B**), and this is a considerably more dramatic loss of LD colocalization than observed for full-length (1-777; **Figure 2C and 2B**) or N-terminally truncated (112-777; **Figures S2D and S2H**) DFCP1 constructs. By contrast, those constructs containing the P-loop domain and the ERB (1-553 and 112-553) remain strongly colocalized to LDs during starvation, countering the starvation-induced loss of LD localization seen in constructs containing the FYVE domains (**Figures 2F, 2B, S2E and S2H**). Since constructs containing the P-loop domain but lacking the ERB are diffuse, this suggests the P-loop domain structurally modulates LD localization, but the ERB is required for maintaining an interaction with the LD membrane.

### DFCP1 is a novel GTPase

To date, the significance of the P-loop in DFCP1 has not been determined. This is surprising given that the P-loop domain is a highly conserved feature of DFCP1 found across all species (**Figure S3A**). Additionally, P-loop domains are essential phosphate-binding motifs found in NTPases. In line with this, cursory amino acid sequence analysis of DFCP1 using HHPRED (Zimmermann et al., 2018) revealed that its P-loop bears similarity to the P-loop found in the GIMAP family of GTPases. Interestingly, GIMAPs are a class of GTPases that have been reported to interact with LDs in immune cells (Schwefel et al., 2013), however, aside from the P-loop, DFCP1 bears little sequence or structural similarity to GIMAP proteins. Nonetheless, this prompted us to identify other common GTPase motifs (G-boxes) in DFCP1 (**Figure 3A**), which would indicate that DFCP1’s P-loop domain is a bona fide GTPase domain. All bona fide GTPases contain 5 essential sequence motifs, called G-boxes (G1-G5), that are critical for GTP binding and hydrolysis (Bourne et al., 1991). Among the G-box elements, G1-G4 show high conservation across major GTPase families. Furthermore, the amino acid separation between G-box elements G1 and G4 is also well conserved within the family of GTPases (**Figure S3B**), with Ras superfamily GTPases having the shortest amino acid separation (∼100 residues) while Gαs have the largest separation (∼225 residues). Based on the number of amino acids separating the last residue of G1 (S194) from the last residue of G4 (L295), the DFCP1 GTPase domain appears to be most similar to a Ras superfamily GTPase, albeit the sequence identity within this region to any member of these families is low.

**Figure 3:**
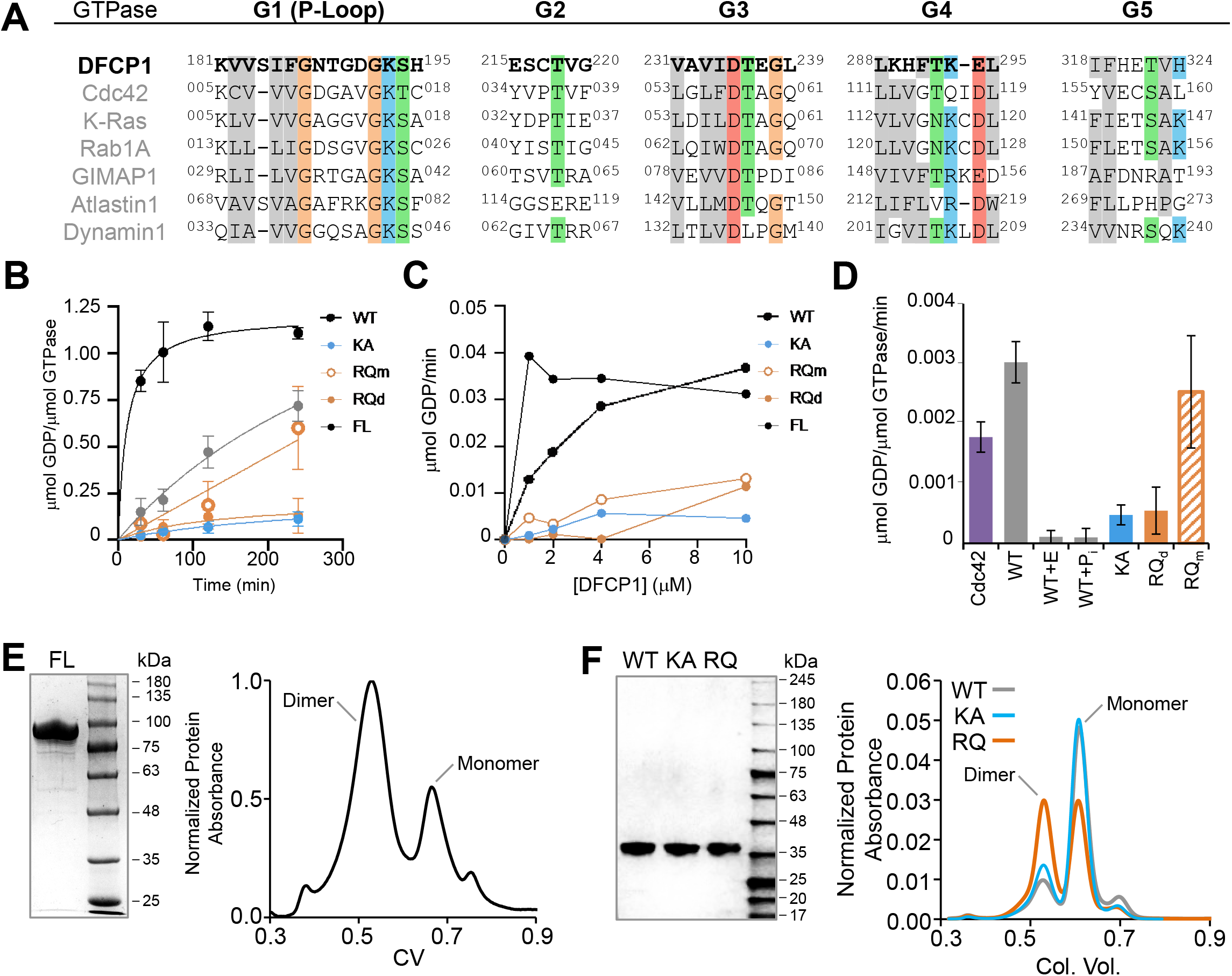
Characterization of the DFCP1 GTPase Domain. **(A)** Sequence alignment of the human DFCP1 G-box motifs with the G-boxes from several representative GTPases. **(B-C)** GDP release assay using 20 μM GTP of the following DFCP1 constructs: FL (residues 1-177; black), WT (112-415; gray), KA (112-415 containing the K193A mutation; blue), or RQ_m_/RQ_d_ (monomeric/dimeric states of 112-415 containing the R266Q mutation; orange open and closed), showing the amount of GDP released over time using 10 μM of the DFCP1 constructs **(B)** and the rate of GDP released as a function of the concentration **(C)** Plotted data points represent mean ±SD for 3 independent experiments. **(D)** Comparison of the specific GDP release rates for 10 μM Cdc42 and the constructs in **Figure 3B** and **3C**. In the case of WT+E and WT+P_i_, 30Mm EDTA or 50Mm phosphate buffer, Ph 7.4 was included in the assay, respectively. Bar graphs represent mean ±SD for 3 independent experiments. **(E)** SDS-PAGE gel and size exclusion chromatography profile of full-length FLAG-DFCP1. **(f)** SDS-PAGE gel and size exclusion chromatography profile of WT, KA and RQ MBP-DFCP1 112-415. See also **Figure S3**.

To confirm that DFCP1 possesses a bona fide GTPase domain, we determined the amount of GDP released by DFCP1 over time (**Figure 3B and S3C**) by measuring the competitive displacement of a fluorescent GDP molecule from a fluorescence-quenching GDP antibody. Using this assay, we found that full-length DFCP1 (FL), purified from mammalian cells, had a robust turnover rate of ∼0.03 μmol GDP μmol GTPase^-1^ min^-1^, which is similar to the basal rate of GTP hydrolysis by Dynamin I (∼0.04 μmol GDP per μmol GTPase per min) measured using the same assay (Mohanakrishnan et al., 2017). By contrast, the isolated DFCP1 GTPase domain (WT) had considerably attenuated GTP turnover rate, ∼0.004 μmol GDP per μmol GTPase^-^ per min. Importantly, the GTPase activity of this domain could be abolished by mutating the highly conserved lysine in the P-loop (K193) to an alanine (**Figure 3B-D**), which is commonly done to disrupt nucleotide binding in bona fide GTPases (Sigal et al., 1986). This isolated GTPase shares many properties with Ras superfamily GTPases, such as a low basal turnover rate that is comparable to a canonical Ras GTPase, such as Cdc42 (**Figure 3D**) and the catalytic activity of the GTPase can be blocked by an excess of magnesium chelating agents such as EDTA, or through a high concentration of phosphate (**Figure 3D**). The difference in the GTPase rates between the full-length protein and the isolated GTPase domain suggests that additional domains in DFCP1 may modulate its own GTPase activity, although it remains unclear which other domains may be responsible.

The Ras-like activity of the DFCP1 GTPase domain is surprising given that the overall domain organization of DFCP1 protein is most similar to a large GTPase, such as Dynamin or Atlastin, which have one or more specialized membrane-binding modules flanking the GTPase domain. Furthermore, these GTPases typically form dimeric or tetrameric complexes, in a nucleotide-dependent manner, which helps to enhance the membrane-binding potential of these GTPases as well as their activities on membrane surfaces (Ford et al., 2011; Moss et al., 2011). Given the architectural similarity of full length DFCP1 with large GTPase, we set out to determine whether DFCP1 forms oligomers under different nucleotide states using size exclusion chromatography (SEC). Using this approach, we found that both full length DFCP1 and the isolated GTPase domain form both monomeric and dimeric states, with full length protein favoring the dimeric state (**Figure 3E and 3F**) and the isolated DFCP1 GTPase domain favoring the monomeric state (**Figure 3F**).

To further probe the relationship between nucleotide activity and dimerization, we introduced a previously unstudied cancer-related mutation into DFCP1. Among all the residues of DFCP1, only the R266Q mutation has been found prevalently in patients (**Figure S3D**). Curiously, this mutation was found to occur in patients with either endometrial and colorectal cancers, which are two cancers distinguished by the presence of an abnormal abundance of LDs (Cotte et al., 2018). We therefore wondered if this mutation could serve as a vital link between DFCP1’s biochemical properties with its role on LDs. Introducing this mutation in the GTPase domain led to the formation of a stable dimeric population of the DFCP1 GTPase domain, which we observed using SEC (**Figure 3F**). What is more, these two oligomeric states show strikingly different GTPase activities: the monomeric R266Q GTPase domain has a turnover rate very similar to the WT GTPase domain, whereas the dimeric R266Q GTPase domain is enzymatically inactive (**Figure 3D**). This mutation suggests that DFCP1 may function like a number of large GTPases, where GTP binding and/or hydrolysis is needed to induce dimerization. However, we were unable to directly modulate such changes in conjunction with non-hydrolysable nucleotide analogs (GTPγS or GMPPNP) or transition state mimics (GDP.VO4- or GDP.Be.X) (data not shown).

### DFCP1’s GTPase activity switches its localization from LDs to autophagosomes

To determine the impact of the DFCP1 GTPase domain on LD metabolism, we rescued Hep3B DFCP1 KD cells by transiently overexpressing full-length GFP-DFCP1 constructs bearing GTPase domain mutations K193A and R266Q **(Figures 4A-C)**. Like in OA treated U2OS cells, WT GFP-DFCP1 uniformly coats LDs in fed and starved OA treated Hep3B cells (**Figures 4A and 4D**). In contrast, GFP-DFCP1 K193A localizes poorly to LDs in both fed or starved cells, and instead forms puncta that remain well-separated from LDs (**Figures 4B and 4D**). The R266Q mutation, on the other hand, shows a similar accumulation on LDs as WT DFCP1 under fed conditions, but this accumulation is markedly increased upon starvation (**Figures 4C and 4D**). Notably, these differences in DFCP1 accumulation on LDs are not a result of expression differences in these constructs (**Figure 4E**).

**Figure 4:**
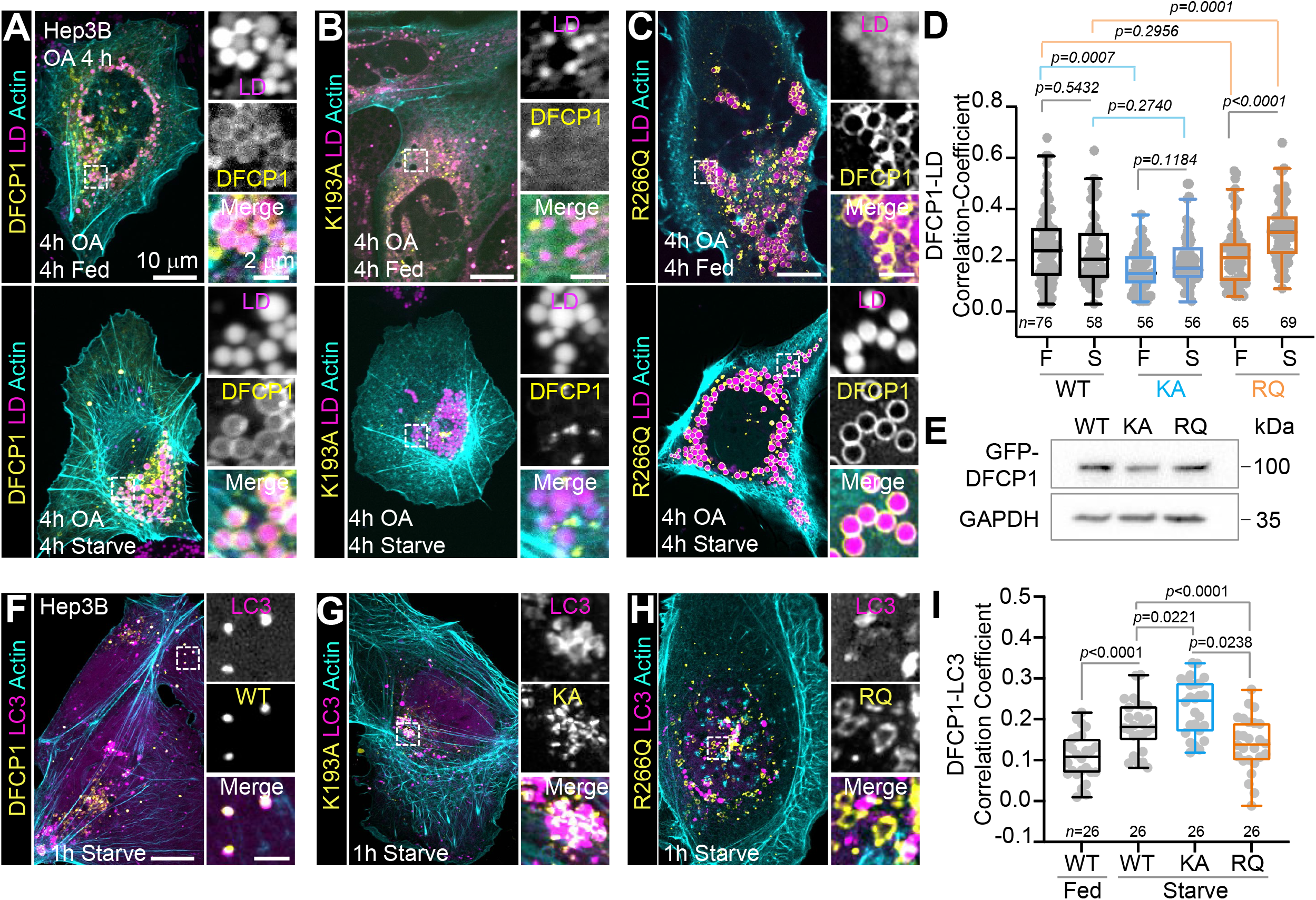
DFCP1 Mutations Modulate LD Autophagosome Localization. **(A-C)** Live-cell confocal images of DFCP1 KD Hep3B cells rescued with either WT (WT, **A**), K193A (KA, **B**), or R266Q (RQ, **C**) GFP-DFCP1 constructs, and stained with LipoTox DR. Prior to imaging, cells were OA-stimulated for 4 h before incubating in either growth or starvation media for 4 h. **(D)** The extent of colocalization (Pearson’s correlation coefficient) between GFP-DFCP1 mutants with LDs from cell populations depicted in **Figure 4A-C**. **(E)** Western blots of cell lysates from the rescued Hep3B cells depicted in **Figure 4A-C**, showing the expression levels of the GFP-DFCP1 mutants after starvation. **(F-H)**. Live-cell confocal images DFCP1 KD Hep3B cells expressing LifeAct-mTagBFP2, mCherry-LC3 and either WT (left), KA (middle), or RQ (right) GFP-DFCP1 constructs. Prior to imaging, cells were incubated in either growth or starvation media for 1 h. **(I)** The extent of colocalization (Pearson’s correlation coefficient) between GFP-DFCP1 and mCherry-LC3 for the cell populations depicted in **Figure 4F-H**. The statistical significance of the measurements was determined using the Mann–Whitney U-test based on the number of observations indicated in each figure panel, which were recorded from two independent transfections. Exact *p*-values are reported. The scale bars in whole-cell and inset images represent 10 and 2 μm, respectively.

As stated above, in the absence of LDs and under conditions of macroautophagy, DFCP1 mobilizes to sites of autophagosome biogenesis on the ER (**Figure S1A**). We therefore wanted to assess whether mutations in the DFCP1 GTPase domain could also modulate the ability of DFCP1 to colocalize with autophagosomes marked by LC3 in starved and rescued Hep3B cells. As previously observed for HEK293 cells (Axe et al., 2008), starvation in the absence of LD stimulation leads to an increase in GFP-DFCP1 puncta that colocalize preferentially with mCherry-LC3 puncta in starved Hep3B and U2OS cells (**Figures 4F and 4I**). This colocalization was further enhanced with the K193A mutation, which mainly forms small puncta that decorate LC3 clusters (**Figures 4G and 4I**). In contrast, the R266Q mutation leads to the formation of ring-like tubular structures, which were well separated, from LC3 puncta (**Figures 4H and 4I**). When taken together, these experiments suggest that the localization to LDs and autophagosomes are anticorrelated and dictated by the DFCP1’s GTPase activity.

### DFCP1’s GTPase activity regulates LD metabolism

Our knockdown experiments indicate that DFCP1 directly modulates LD metabolism (**Figure 1G-I**), and given that the DFCP1 GTPase domain is important for its localization to LDs (**Figure 2 and Figure 4)**, then GTPase mutations could modulate the LD metabolism (**Figure 5**). Under fed conditions, Hep3B DFCP1 KD cells rescued with either GTPase mutations are able to generate equivalent numbers of LDs (**Figure 5A**) as WT rescued cells, but the size distribution of these LDs differ significantly between each construct (**Figure 5B**). Rescuing with the K193A mutant causes LDs to be significantly smaller than those found in the WT rescue cells, whereas rescuing with the R266Q mutant causes LDs to be marginally larger than those found in WT rescue cells. Upon starvation, the number of LDs in WT and K193A rescued cells increases, with the number of LDs in the starved K193A rescue cells being significantly more than the number formed in WT. This change in LD number is anticorrelated with the size of LDs, as the K193A rescue cells have LDs that are considerably smaller than those found in WT rescue cells. In contrast to WT and K193A, cells rescued with R266Q mutation do not show a significant change in the density of LDs, but have LDs that are considerably larger than those found in WT cells. This suggests that the LD size distribution depends on the GTPase activity of DFCP1.

**Figure 5:**
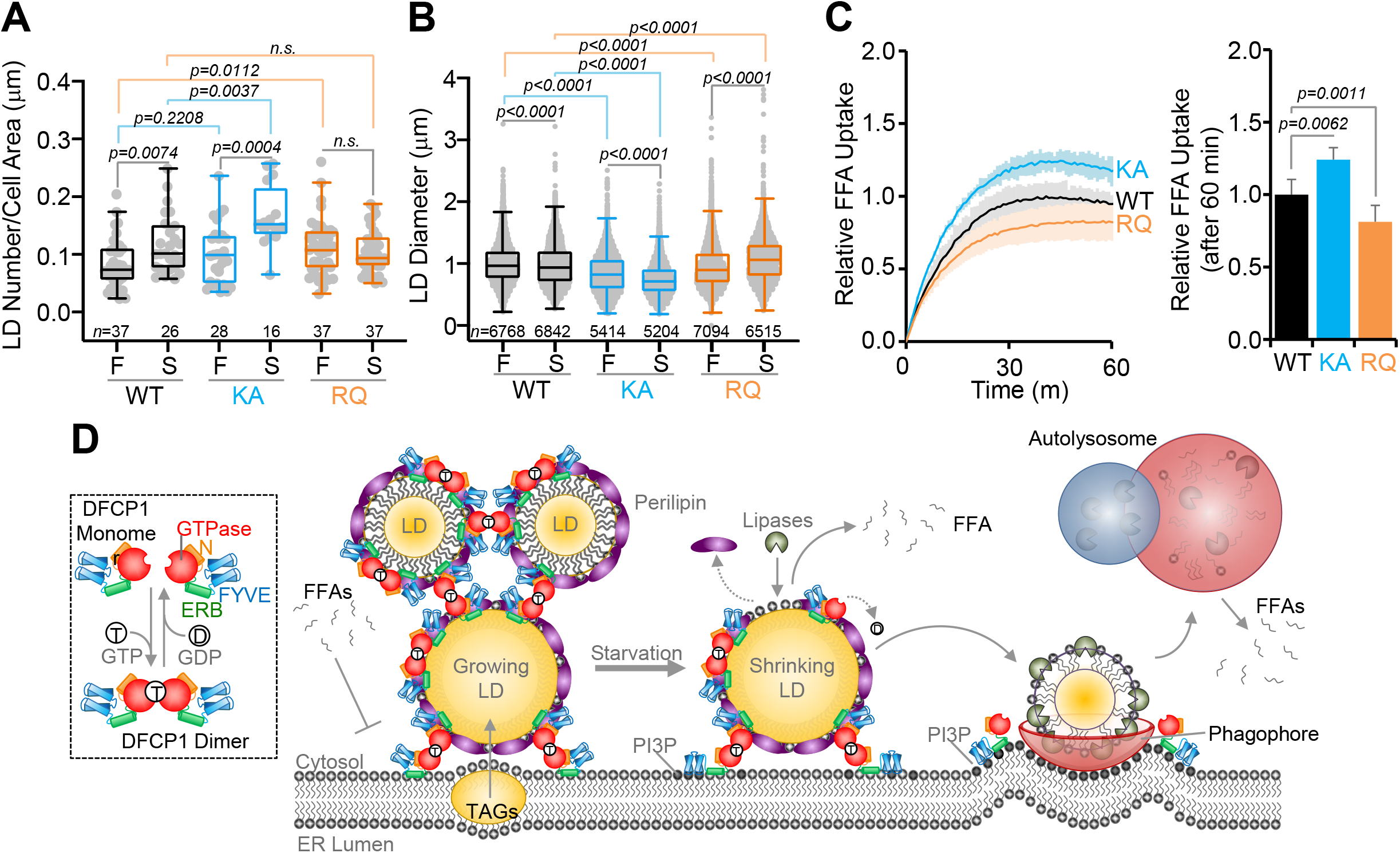
DFCP1 Mutations Modulate Fatty Acid Storage. **(A,B)** Density and diameter distributions of LDs quantified from images represented in **Figure 4A-C**. LD density was determined by dividing the total number of LDs for a cell by the cell’s area. LD diameter was measured in in the plane where a given LD’s diameter was the largest. LDs were visualized using LipoTox DR. **(C)** FFA uptake measured in KD Hep3B cells cultured under normal growth conditions and rescued with WT, KA or RQ GFP-DFCP1 constructs, as well as LifeAct-mTagBFP2. **(D)** Model of the role of DFCP1’s GTPase domain in LD metabolism. The GTP binding to DFCP1 facilitates oligomerization and localization of DFCP1 to LDs. DFCP1 preferentially accumulates on LDs in fed cells where it ultimately impairs fatty acid uptake and inhibits the breakdown of LDs. Upon starvation, DFCP1 translocates from LDs to autophagosomes to facilitate lipophagy. The statistical significance of the measurements was determined using the Mann–Whitney U-test based on the number of observations indicated in each figure panel, which were recorded from two independent transfections. Exact p-values are reported with exception to p>0.05, which are considered to be nonsignificant (n.s.).

This pattern of DFCP1 localization further correlates with changes in FFA uptake (**Figure 5C**). DFCP1 KD Hep3B cells rescued with endogenous levels of DFCP1 K193A are able to take up ∼25% more FFAs than cells rescued with WT DFCP1, recapitulating the behavior observed in DFCP1 KD cells rescued with GFP (**Fig. 1H**). By contrast, those cells rescued with DFCP1 R266Q show ∼25% reduction in FFA uptake. This suggests that DFCP1 accumulation on LDs indirectly regulates fatty acid flux into cells. When taken together, our data indicates that DFCP1 accumulation on LDs is directly regulated by its GTPase activity, and this accumulation impedes both LD clearance and FFA uptake. Thus, DFCP1 functions as a molecular switch to gate the catabolism in response to nutrient availability (**Figure 5D**).

## Discussion

LDs are transient energy storage depots that help to buffer cellular energy demands. As a consequence, LDs can rapidly form when FAs are abundant but are also quickly broken down when ATP becomes scarce. While the physiological roles of LDs are well established, the molecular regulation of LDs remains poorly understood. In this study, we expand on emerging reports that the autophagy-associated protein DFCP1 is a regulator of LD metabolism (**Figure 5D**). Specifically, we show that DFCP1 accumulation impairs both FA uptake and LD breakdown during nutrient stress (**Figure 1**). This accumulation on LDs is controlled by a previously uncharacterized GTPase domain that works in conjunction with ER-binding motif to regulate its association to the periphery of LDs (**Figure 2**). This GTPase domain is unique in that it possesses a low GTP turnover rate commonly found in small Ras-like GTPases, but also has the ability to oligomerize in a GTP-dependent manner that is commonly found in large GTPases (**Figure 3**). In cells, modulating the function of this GTPase domain through mutagenesis leads to aberrant accumulation of DFCP1 on LDs and autophagosomes (**Figure 4**). This change in DFCP1 localization is also coupled to a change in LD size distribution and FA metabolism (**Figure 5**). Thus, DFCP1 represents a novel GTP-dependent molecular switch that dictates the availability of LDs for energy consumption by the cell (**Figure 5D**).

Large oligomeric membrane-associating GTPases, like DFCP1, are well known to play a direct role in the fission and fusion of cellular organelles, including: the plasma membrane, early, late and recycling endosomes, the Golgi, mitochondria, and the ER. For LDs, the role of such GTPases has been less studied. To date, association of large GTPases with LDs has only been observed in lymphocytes, which uniquely possess a family of proteins called the GTPases of Immunity Associated Proteins (GIMAPs) make up a family of septin-like proteins have been shown to regulate LD metabolism (Limoges et al., 2021). For example, GIMAP2 has been found to localize directly to LDs, using its dual hydrophobic C-terminal stretches to associate and potentially tether LDs to each other (Schwefel et al., 2010). Consequently, GIMAP2 overexpression in these cells causes LD numbers to increase. Blocking the ability of GIMAP2 to oligomerize abolishes this expression-induced increase in LDs. GIMAPs have not been shown to regulate LDs outside lymphocytes, and thus, it appears that the universally expressed DFCP1 may replace GIMAPs in other cell types. Interestingly, DFCP1 shares a P-loop sequence that most closely resembles the P-loop in GIMAP GTPases **(Figure 3A)**, although catalytic GTPase domain that mostly resembles a Ras-like GTPase. The latter maybe important in understanding why the expression of GIMAPs and DFCP1 may have opposite effects on LD accumulation.

Large GTPases are also known to be involved in modulating contacts between the ER and other organelles. For example, Mitofuisn 2 tethers and regulates the spacing between the outer Mitochondrial membrane to the ER (Filadi et al., 2015). Similarly, the dynamin-like Atlastins were shown to regulate tethering of Cop II coated vesicles to the ER (Niu et al., 2019). In both of these cases, tethering is driven by GTP-dependent oligomerization of the GTPase domain and multi-valent membrane binding to the ER and another organelle. While DFCP1 shares the same basic molecular architecture, including the ability to dimerize in a GTP-dependent manner and the capacity to simultaneously bind to multiple membrane compartments, DFCP1 was only shown to establish an ER-LD contact site through the formation of a tethering complex consisting of NAG-RINT1-ZW10 (NRZ) and Rab18 (Li et al., 2019). In so doing, this complex is able to bring LDs in close proximity to the ER in order to drive LD expansion during biogenesis (Li et al., 2019). During macroautophagy, the NRZ complex and it’s components are known to take on more passive roles in autophagy, such as tethering components needed for autophagy initiation at the ER or the mobilization of ATG9 out of the TGN (He et al., 2013) By contrast, DFCP1 has been shown to accumulate on PI3P-rich autophagosome precursor sites on the ER, known as omegasomes (Axe et al., 2008). This suggests that DFCP1 may also function as an autonomous tethering factor in LD metabolism. While the role of DFCP1 in autophagy remains puzzling, it’s possible that PI3P-binding by DFCP1 ensures that LDs are tethered to the ER near sites of autophagosome formation. Additionally, this binding to PI3P could also serve to sequester DFCP1 that has undergone GTP-dependent disassembly from LD surface. In this way, DFCP1 could promote tethering of LDs near degradative subdomains of the ER, which would then be released for rapid catabolism in response to an environmental trigger.

While the impact of DFCP1 on LD metabolism is clear, the specific phase of LD breakdown regulated by DFCP1 remains uncertain. Recently, it has been suggested that LDs follow a two-step catabolic pathway. First, larger LDs are reduced in size through the combined actions of perilipin removal by chaperon mediated autophagy and the breakdown of TAGs housed within the LD by LD-associated lipases (Kaushik and Cuervo, 2015). This process, which is thought to require declustering and/or a separation of LDs from the ER (Pfisterer et al., 2017), ultimately results in LDs with reduced volume. When LDs become sufficiently small, they can be consumed in whole by the canonical autophagy system (Schott et al., 2019). Consequently, impairing lipolysis leads to a net increase in the number of larger LDs, since smaller LDs would be preferentially cleared by the autophagy system. Whereas, impairing autophagy is expected to result in an accumulation of small LDs that have been processed from large LDs undergoing lipolysis (Schott et al., 2019). We have observed that knockdown of DFCP1 leads to changes in the LD size distribution that is consistent with inhibition of autophagy and suggests that DFCP1 is needed for autophagic clearance of small LDs (**Figures 1G and 1H**). However, knockdown of DFCP1 does not directly impair autophagosome formation (**Figure S1B**), which suggests the role of DFCP1 may be upstream of lipophagy. One possibility is that DFCP1 may slow down the rate of lipolysis, and thereby give more time for the autophagy system to clear the small LDs and prevent their accumulation. However, this assumption would require lipophagy-dependent clearance of LDs to be slower than the rate by which lipolysis reduces the size of LDs. At this time the relative contributions of these pathways on the dynamics of LD metabolism is not known and thus both modes of regulation remain possible.

In summary, we have shown that DFCP1 is a novel GTPase that accumulates on LDs under conditions that promote LD biogenesis in order to impair LD catabolism. However, during macroautophagy, DFCP1 redistributes away from LDs on to the ER to promote LD catabolism. Thus, DFCP1 represents a novel molecular switch that is capable of promoting both LD growth and LD catabolism.

## Supporting information

Supplemental Manuscript

## Supplemental Information

Supplemental information includes experimental methods and three figures.

## Acknowledgments

This work was supported by grants from the National Institutes of Health (R01 GM136925) and the Elsa U Pardee Foundation (P20-03924). We would also like to thank Dr. Silvia Jansen for her critical reading of the manuscript.

