## Supplemental Manuscript for "The GTPase Activity of the Double FYVE Domain Containing Protein 1 (DFCP1) Regulates Lipid Droplet Metabolism"

### Methods

**Mammalian Cell Culture:** Hep3B and U2OS cells were cultured at a 37 °C with 5% $CO_2$ and in growth media consisting of MEM GlutaMax or DMEM GlutaMax supplemented with 10% FBS and antibiotic-antimycotic (Thermo Fisher Scientific, Waltham, MA), respectively. Two days prior to live-cell imaging experiments, 50,000 cells were seeded on 3.5 cm imaging dishes and grown to 40% confluency prior to transfection with the indicated plasmids using FuGENE HD (Promega, Madison, WI). In some cases, cells were also incubated for 2 days before imaging with a siRNA mixture, consisting of 5 μL of RNAiMax (ThermoFisher Scientific) and 2.5 pmol of the specified siRNA. Cells were induced to form LDs by supplementing the growth media with 200 μM oleic acid for 4 or 24 h (as indicated), before exchanging the media with either normal growth media or EBSS for 1, 4, or 18 h (as indicated) prior to imaging. LDs were identified by incubating with the lipid droplet specific dyes Bodipy-C12-568 (ThermoFisher Scientific), LipoTox Green (ThermoFisher Scientific) or LipoTox Deep Red (ThermoFisher Scientific) at volume ration of 1:10,000 for 30 min prior to imaging. In some cases (**Figure 1A**), cells were treated with 1 μM Wortmannin (Cayman Chemical Company, Ann Arbor, MI) for 30 min prior to imaging.

**Live Cell Imaging and Image Analysis:** All cells were imaged using either a Nikon Ti2 inverted microscope equipped with a 100× (1.4 NA) Plan-Apo oil immersion objective and a Yokogawa CSU-W1 spinning disk confocal attached to a Hamamatsu ORCA-FLASH4.0 CMOS camera. Cells in either DMEM FluoroBrite (ThermoFisher Scientific) medium supplemented with 5% FBS or EBSS lacking phenol red (MilliporeSigma) were imaged at 37 °C and 5% $CO_2$. Images stacks were captured at 16-bit 2048 x 2044 resolution with an axial spacing of 0.2 μm using the Nikon Elements Software package. All images were captured blindly and randomly, which involved imaging the nearest cell to a random set of x-y coordinates that contained actin fluorescence (depending on the experiment). Captured images were blinded again and image analysis was performed using the software Fiji (https://imagej.net/Fiji). Specifically, lipid droplet number and diameters were scored manually and 2D/3D colocalization analysis was performed using the Fiji Coloc2 analysis tool. Specifically, LD diameter was measured in the plane where a given LD's diameter was the largest and lipid droplet density was determined by dividing the total number of LDs by the area of the cell's basement membrane.

**GDP Release Assay**: All GTPase assays were performed with the GDP-FI Transcreener assay (Bellbrook Labs, Fictchburg, WI) according to the manufacturer's suggested protocol. In brief, GTPase constructs was added to a tube containing GDP Release Assay buffer (3 mM $MgCl_2$, 1mM DTT, 150mM NaCl, 20mM Tris pH 8) and 20 μM GTP and incubated at RT in 25 μL reactions. Reactions were quenched by the addition of 25 μL stop solution (18.8 μg/mL Ab-IR dye QC-1, GDP-AlexaFluor 594 tracer, and Stop and Detect solution) and immediately read using a platereader for 1h, taking measurements every ten minutes.

**FFA Uptake and Content Assays:** To assess free fatty acid uptake, DFCP1 KD Hep3B cells rescued with the indicated DFCP1 constructs and seeded in a 96-well poly-lysine coated tissue culture plate. Cells were serum deprived for 1 h before adding the fluorescent TF2-C12 Fatty Acid (MilliporeSigma, Burlington, MA). Fluorescence emission was monitored for 1 h at 37 °C on a Citation 5 microplate reader (BioTek, Winooski, VT), with the excitation monochromator set to 485 ±7 nm and the emission monochromator set to 515 ±7 nm. Free fatty acid concentrations from Hep3B cells were measured with Free Fatty Acid Quantitation Kit reagents according to the manufacturers instructions (MilliporeSigma).

**Lipid Droplet Isolation**: Hep3B cells were grown to confluence in 15 cm dishes and treated with 200 μM oleic acid for 16 h to promote the production of lipid droplets. The cells were then washed twice with ice-cold PBS and harvested using a cell scraper. Cells were pelleted at 250 × *g* for 10 min at 4 °C and resuspended in 5 pellet volumes of ice-cold lysis buffer consisting of 20mM Tris, pH7.4, 1mM EDTA. The cell suspension was incubated on ice for 10 min and then lysed by pipetting through a 27-gauge needle 5 times. Cell lysate was centrifuged for 10 min at 1000 × *g* at 4 °C to remove nuclei and insoluble cellular aggregates. The supernatant was collected and mixed with ice-cold lysis buffer with 20% sucrose. Cell lysates were then added to an ultracentrifuge tube and 5mL ice-cold lysis buffer with 5% sucrose was layered gently over the sample. A layer of ice-cold lysis buffer was added over the sucrose layers to fill the tube. The solution was centrifuged for 1 hour at 28,000 × *g* at 4 °C. The floating LD layer was collected and placed in a microfuge tube, which was then centrifuged for 10 min at 20,000 × *g* at 4 °C. The bottom soluble fraction was removed, and the process was repeated until the LD layer was reduced to a final volume of 50 μL.

**Western Blotting**: Cells for western blotting were treated the same way as those for imaging, except, in this case, 200,000 cells were seeded onto a 6 cm plate and the transfection mixtures were doubled in volume. Cells were harvested 3 days post-seeding in lysis buffer consisting of TBS supplemented with 1% Triton-X, 2 mM EDTA with 1 mM PMSF. The lysis mixture was pipette mixed and incubated on ice for 30 min. The lysate was then clarified at 12,000 × *g* for 10 min and the supernatant was removed was mixed with 3X SDS sample buffer. Samples were run on 12% polyacrylamide gels and transferred onto Immobilon-P polyvinylidene difluoride (PVDF) membranes (MilliporeSigma) membrane in 1X transfer buffer with 10% ethanol and 0.01% SDS at 130 mA for 3 h at 4 °C. Membranes were allowed to dry and then blocked for 1 h with 5% BSA in TBS supplemented with 0.1% Tween (TBS-T), before incubating with the specified primary antibodies for 4 h to overnight at 4 °C. Blots were then washed with TBS-T and incubated with horseradish peroxidase (HRP)-linked anti-mouse and anti-rabbit immunoglobulin G secondary antibodies for 30 min at room temperature. Washed blots were developed using the clarity western ECL substrate (Bio-Rad Laboratories, Hercules, CA) and imaged using a Gel-Doc Imager (Bio-Rad). The relative abundance of proteins was determined by densitometry analysis using the program Fiji.

**Protein Expression and Purification.**

*FL hsDFCP1.* Human DFCP1 (Uniprot ID: Q9HBF4) was cloned into a custom vector, where the DNA sequence encoding GFP in a pEGFP-C1 vector was replaced with a DNA sequence encoding a FLAG-TEV sequence using the NheI and BglII restriction sites. For expression of FLAG-DFCP1, $1.25 \times 10^6$ Expi293 cells $mL^{-1}$ (ThermoFisher Scientific), cultured in Expi293 media (ThermoFisher Scientific) without antibiotic and antimyotics, were transiently transfected with a solution of 1 μg $mL^{-1}$ plasmid DNA and 1 mg $mL^{-1}$ polyethylenimine (Polysciences,

Warrington, PA). After 48 h expression, cells were harvested, pelleted, and resuspended in lysis buffer comprised of 25 mM Tris, pH 8.0, 300 mM NaCl, 5% glycerol, 1% Triton-X, 1 mM PMSF, 1X protease inhibitor tablet without EDTA (ThermoFisher Scientific). Cells were incubated in lysis buffer on ice for 30 min and insoluble cellular components were removed by centrifugation at 12,000 × *g* for 10 min. Clarified lysates were incubated with 1 mL Anti-DYKDDDDK (FLAG) Affinity Resin (Genscript; Piscataway, NJ) for 2 h at 4 °C and then washed with 15 column volumes (CVs) of wash buffer, comprised of 25 mM Tris, pH 8.0, 300 mM NaCl, 5% glycerol, and 1 mM PMSF, followed by 15 CVs of wash buffer supplemented with 1 mM ATP. Bound DFCP1 was eluted by 4 sequential 1 h incubations of the FLAG resin with 1 CV of wash buffer supplemented with 1 mg $mL^{-1}$ 3X-FLAG peptide. The elution fractions were pooled, and the excess FLAG peptide was removed by dialysis using a 3-14 kDa cutoff, with a dialysis buffer composed of 25 mM Tris, pH 8, 150 mM NaCl, 1 mM DTT, 1 mM $MgCl_2$. The dialyzed protein was spin concentrated using a 30 kDa MWCO ultracentrifugation filter (MilliporeSigma) and subjected to size exclusion chromatography using a Superose 6 attached to an AKTA Pure 25L (Cytiva Life Sciences, Marlborough, MA) equilibrated in a buffer consisting of 25 mM Tris pH 8, 150 mM NaCl, 1 mM DTT, and 1 mM $MgCl_2$. The resulting monomer and dimer peaks were isolated together, spin concentrated using a 30 kDa MWCO ultracentrifugation filter, and flash frozen in liquid nitrogen.

*DFCP1 GTPase domain and GTPase domain mutations.* Residues 112-415 of mouse DFCP1 (Uniprot ID: Q810J8), which contains a GTPase domain that is 98.8% identical to human DFCP1, was inserted into the pMal-c2e bacterial expression vector using the EcoRI and Sal1 restriction enzyme cleavage sites. DFCP1 GTPase mutations K193A and R266Q were introduced using the QuikChange mutagenesis kit (Agilent, Santa Clara, CA). These constructs were transformed into *Rosetta* (DE3) cells (Novagen) that were subsequently cultured in Terrific Broth medium that was supplemented with 100 μg $mL^{-1}$ carbenicillin and 100 μg $mL^{-1}$ chloramphenicol at 37 °C. Protein expression was induced with 1 mM IPTG at 18 °C for 16 hr. Cells were homogenized in lysis buffer, comprised of 25 mM Tris pH 8.0, 300 mM NaCl, 1 mM $MgCl_2$, 4 mM benzamidine hydrochloride, and 1 mM PMSF, and lysed using a microfluidizer 110L (Microfluidics, Newton, MA). Lysates were clarified by centrifugation at 16,000 × *g* for 45 min and the supernatant was loaded onto amylose affinity resin (New England BioLabs, Ipswich, MA). The column was washed with 10 CVs of lysis buffer and the protein was eluted with sequentially added 1 CV aliquots of lysis buffer supplemented with 20 mM maltose. Aliquots containing the eluted protein were combined and passed over a HiLoad 26/600 Superdex 200 pg size exclusion column (Cytiva) equilibrated in 25 mM Tris pH 8.0, 150 mM NaCl, 1 mM $MgCl_2$, and 1 mM DTT. The monomer and dimer peaks were isolated and the protein was spin concentrated to 2 mL volume using a 10 MWCO ultracentrifugation filter. The MBP tag was removed by treating the concentrated protein solution with 0.15 mg $mL^{-1}$ TEV protease overnight at 4 °C. The TEV-cleaved protein was filtered and flashed diluted with 2 volumes of dialysis buffer lacking NaCl in order to reduce the NaCl concentration to 50 mM. The cleaved protein was further purified with a MonoQ 4.6/100 PE column (Cytiva), using a 50 mM to 300 mM NaCl gradient spanning 25 CVs. The peak corresponding to DFCP1 was isolated, spin concentrated, and then flash frozen in liquid nitrogen.

*Cdc42.* WT human cdc42 (Uniprot ID: P60953) was inserted into the pTYB11 IMPACT bacterial expression vector (New England BioLabs), using the Sal1 and SapI restriction enzyme cleave sites. Cdc42 was transformed into *Rosetta* (DE3) cells (Novagen) that were subsequently cultured in Terrific Broth medium, supplemented with 100 μg $mL^{-1}$ carbenicillin at 37 °C. Protein expression was induced with 1 mM IPTG at 18 °C for 16 hr. Cells were homogenized in lysis buffer comprised of 25 mM Tris pH 8.0, 300 mM NaCl, 5 mM $MgCl_2$, 0.1 mM GDP, 4 mM benzamidine hydrochloride, and 1 mM PMSF, and lysed using a microfluidizer 110L (Microfluidics). Lysates were clarified by centrifugation at 16,000 × *g* for 45 min and the supernatant was loaded onto the chitin affinity resin (New England BioLabs). The affinity tag was removed by inducing self-cleavage of the intein domain with 50 mM DTT for 48 h at 4 °C. Proteins were eluted from the column in 25 mM Tris pH 8.0, 300 mM NaCl, 5 mM $MgCl_2$, 0.1 mM GDP and 50 mM DTT. A final size exclusion chromatography purification step was performed on a HiLoad 26/600 Superdex 75 pg column (Cytiva) equilibrated in 25 mM Tris pH 8.0, 50 mM NaCl, 2 mM $MgCl_2$, 0.1 mM GDP and 1 mM DTT. The protein was spin concentrated using an ultracentrifugation filter and flash frozen in liquid nitrogen.

### Supplemental Figures:

#### Figure S1: DFCP1 Regulates LD Metabolism.

**(A)** Live cell confocal images of U2OS cells expressing BFP-DFCP1, GFP-LC3 and mCherry-Sec61β. Cells were incubated in either growth (left) or starvation media (middle). The extent of colocalization (Pearson's correlation coefficient) between BFP-DFCP1 and GFP-LC3 is plotted on the right.
**(B)** Western blot showing the conversion of endogenous LC3-I to LC3-II in clarified cell lysates from NT and KD U2OS cells that were either fed or starved for 4 h prior to harvesting. Graph shows LC3I/II ratio normalized to GAPDH, as determined by densitometry of blots ($n$=11).
**(C)** Western blots of purified LDs isolated from fed and starved OA-stimulated U2OS cells expressing GFP-DFCP1 showing the total cell lysate (L), the last wash (W), and the purified LD fraction (LD).
**(D)** FFA content measured in NT or KD OA-stimulated Hep3B cells that were either fed or starved for 4 h. Bar graphs represent mean ±SD for 4 independent experiments.
**(E)** Number and diameter distributions of LDs quantified from images of fed and starved untransfected OA-stimulated Hep3B cells. LDs were identified by treating cells with LipoTox DR for 30 min prior to imaging.

The scale bars in whole-cell and inset images represent 10 and 2 μm, respectively.

The statistical significance of the measurements was determined using the Mann–Whitney U-test (**A and E**), Student's t-test (**D**) or the Wilcoxon matched-pairs signed rank test (**B**) based on the indicated number of observations (indicated in the figure panel) recorded from at least two independent transfections. Exact $p$-values are reported with exception to $p$>0.05, which are considered to be nonsignificant (n.s.)

#### Figure S2: Domain Requirements for DFCP1 Localization.

**(A)** Domain diagram of DFCP1 depicting the GFP-tagged constructs used in this figure (**C-G**).
**(B)** Western blot showing expression levels of the all GFP-DFCP1 truncations depicted in **Figure S2A** in U2OS cells.
**(C-G)** Representative images of U2OS cells expressing LifeAct-mTagBFP2 and either GFP-Sec61β or the indicated GFP-DFCP1 truncations. Prior to imaging, all cells were OA-stimulated for 20 h before incubating in either growth (fed) or starvation media (starved) for 18 h. LDs were stained by treating cells with LipoTox DR for 30 min.
**(H)** The extent of colocalization (Pearson's correlation coefficient) between GFP-DFCP1 and LDs from cell populations depicted in **Figures 2C-F** and **Figures S2C-G.** The extent of colocalization between GFP-Sec61B and LDs is also included as a reference.

The scale bars in whole-cell and inset images represent 10 and 2 µm, respectively.

The statistical significance of the measurements was determined using the Mann–Whitney U-test. Exact p-values are reported.

**Figure S3: Identification of the DFCP1 GTPase Domain.**
**(A)** The percent identity across species for the full length human DFCP1 protein (open bars) and the DFCP1 GTPase domain (striped bars), showing that the GTPase domain is highly conserved.
**(B)** The average number of residues separating G-box 1 and G-box 4 in the indicated family of GTPases. Notably, the DFCP1 GTPase domain has a separation similar to that of the Ras superfamily GTPases.
**(C)** GDP release assay showing the amount of GDP over time for 2 μM of the GTPase constructs used in Fig 3. Plotted data points represent mean ±SD for 3 independent experiments.
**(D)** Chart of the frequency of somatic cancer mutations in DFCP1 found in 4440 TCGA tumor samples (https://www.cancer.gov/tcga).
