## Supplemental Figures for "The GTPase Activity of the Double FYVE Domain Containing Protein 1 (DFCP1) Regulates Lipid Droplet Metabolism"

### Supplemental Figure 1: Domain Requirements for DFCP1 Localization

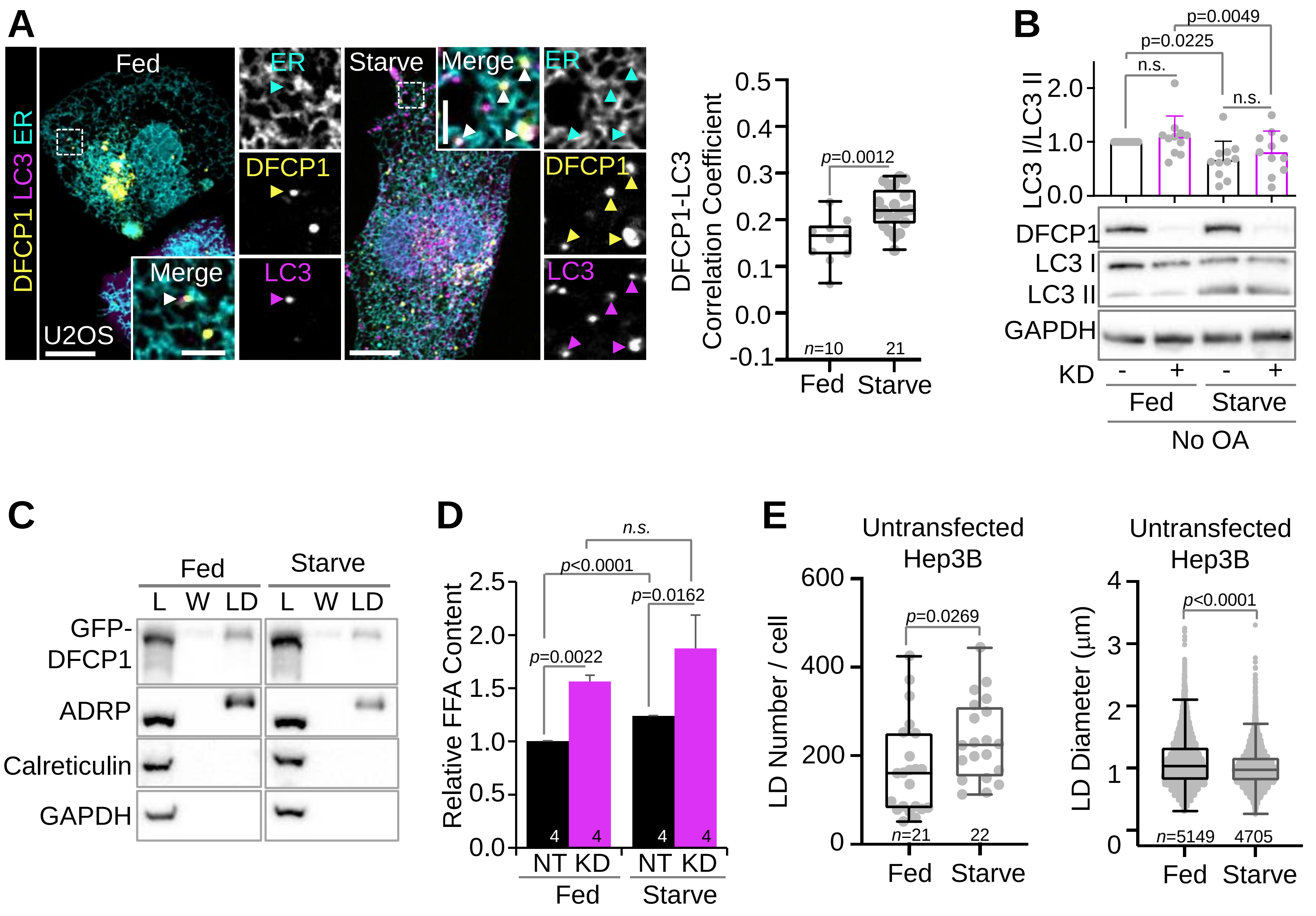

### Supplemental Figure 2: Domain Requirements for DFCP1 Localization

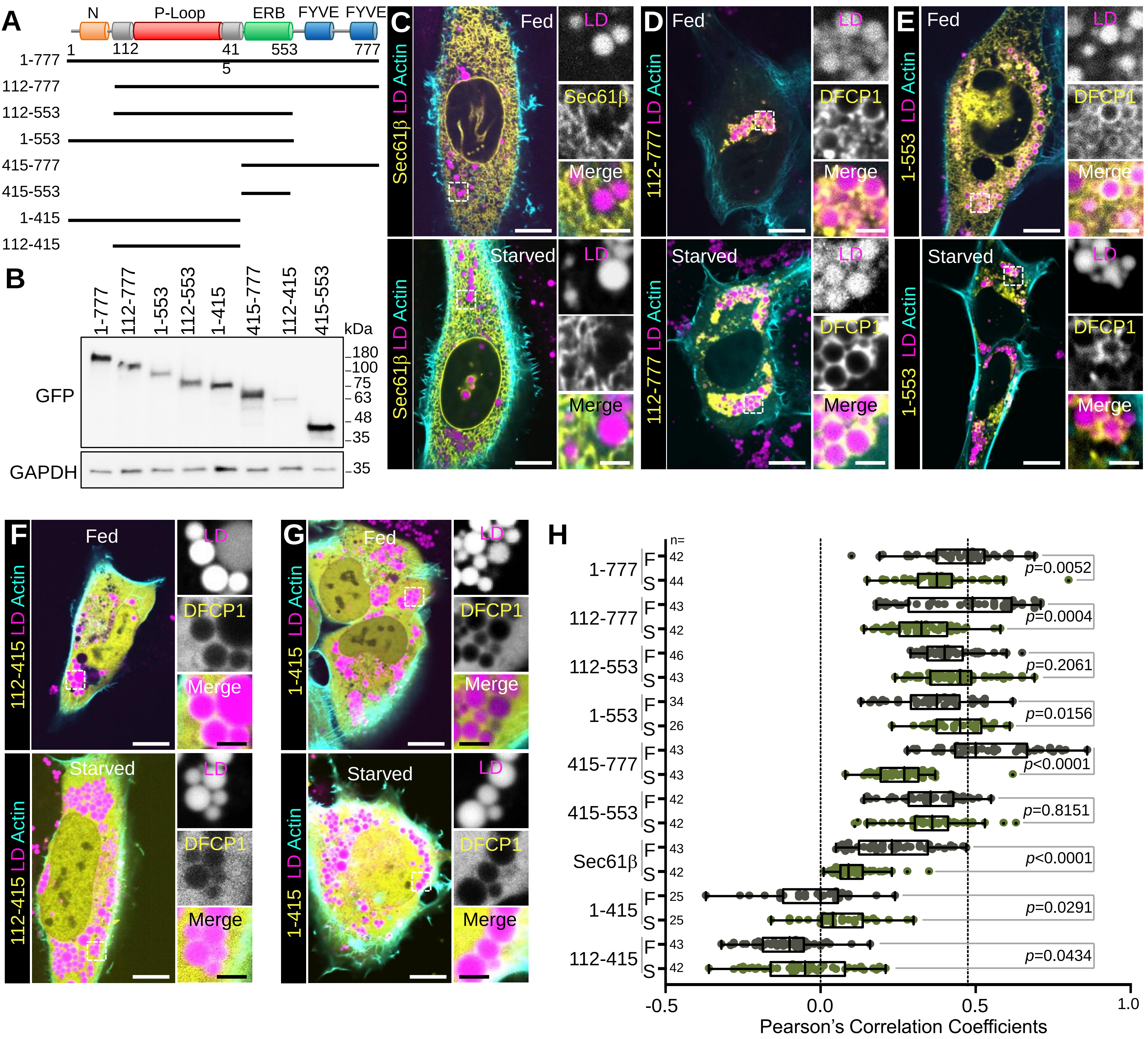

### Supplemental Figure 3: GTPase Activity of DFCP1

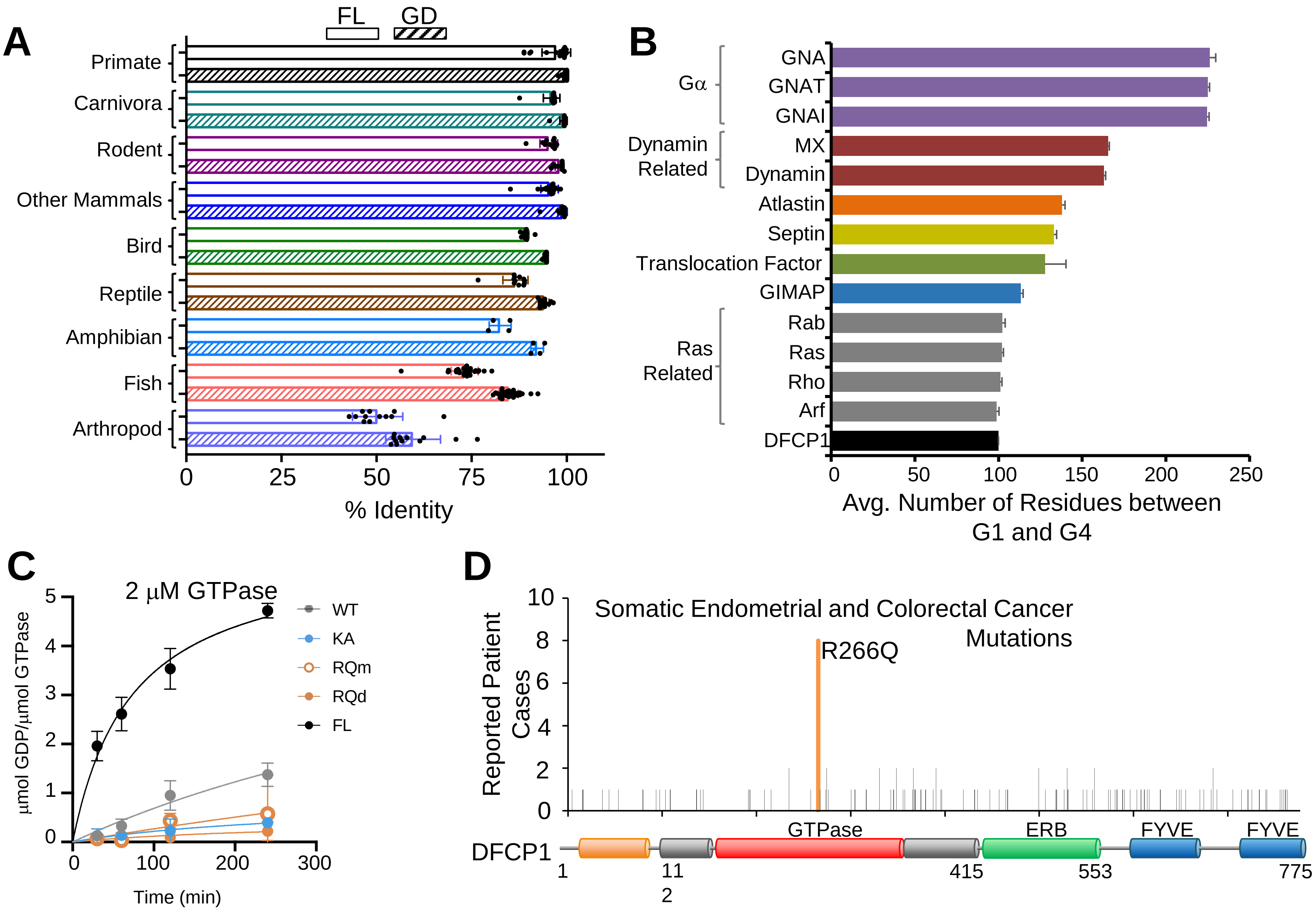
